# Autonomic Decoupling in Listening Effort: Why Single-Channel Biomarkers Fail

**DOI:** 10.64898/2026.07.24.740555

**Authors:** Stefan Bleeck, Yuki Wang, David Simpson

## Abstract

**Objective:** Current theoretical frameworks operationalize listening effort as a monolithic, unified sympathetic response, driving the search for a single universal clinical biomarker (predominantly task-evoked pupillometry). This study stress-tests the “unified arousal” hypothesis by evaluating the continuous cross-modal overlap of autonomic and cortical responses during a speech-in-noise paradigm.

**Design:** Continuous, simultaneous physiological tracking-including Pupillometry (Locus Coeruleus), Galvanic Skin Response (GSR), Cardiac Inter-beat Interval (IBI), Respiration Rate, and Frontal Alpha EEG-was conducted on N=28 normal-hearing adults. Participants completed a continuous speech-in-noise task (OLSA) across varying Signal-to-Noise Ratios (+12 dB to -16 dB). A continuous Spearman’s rank correlation matrix was utilized to assess physiological decoupling, and Linear Mixed-Effects (LME) models isolated single-trial effort dynamics.

**Results:** The data revealed a lack of meaningful cross-modal overlap. Continuous correlation matrices demonstrated negligible relationships across the four autonomic channels (all p>0.05). Trial-level cross-correlations confirmed extremely weak functional coupling across the autonomic nervous system (mean effect sizes strictly bounded between ρ=−0.042 and ρ=0.065). These negligible effect sizes fail to support the “unified sympathetic storm” hypothesis, providing strong evidence for heavy physiological decoupling, where autonomic channels fail to synchronize during acute stress.

Furthermore, subjective retrospective effort was highly collinear with objective task difficulty (r=0.848), offering limited independent tracking of internal state.

**Conclusions:** Listening effort is not a unified whole-body state; it is an idiosyncratic, heavily decoupled biological routing process. The reliance on single-channel tracking is structurally flawed, as it is blind to autonomic diversity. Future models of cognitive exertion must decouple generalized capacity allocation from idiosyncratic autonomic routing, and experimental paradigms must separate genuine effortful limits from active task withdrawal.

## 1. Introduction

Listening effort is currently theorized as a monolithic biological event. Dominant frameworks, such as the Framework for Understanding Effortful Listening (FUEL), conceptualize cognitive resource allocation as a unified sympathetic drive. Under this paradigm, encountering degraded speech triggers a generalized whole-body arousal: the locus coeruleus floods the cortex with norepinephrine, electrodermal activity spikes, and cardiovascular rhythms shift in concert. Effort is treated as the physiological “weather” of the auditory system-a transient, pervasive storm that scales uniformly with task demand.

Driven by this assumption of unified arousal, the search for a clinical biomarker of listening effort has largely relied on single-modality methodologies, predominantly task-evoked pupillometry. If sympathetic arousal is monolithic, sampling a single biological channel should theoretically provide a proxy for global cognitive expenditure. However, this approach yields highly variable inter-subject data and frequently demonstrates near-zero correlations across behavioural, subjective, and physiological metrics (Strand et al., 2018). The premise that a single “exhaust pipe” can accurately index the output of the entire cognitive engine remains unverified.

This conceptualization ignores the possibility of autonomic decoupling. Rather than a uniform physiological response, listening effort may be dictated by stable, idiosyncratic routing preferences, resulting in the functional decoupling of autonomic channels. Confronted with identical acoustic degradation, one listener may default to a massive locus coeruleus response with minimal electrodermal engagement, while another relies entirely on respiratory pacing. If effort is decoupled across independent physiological channels, single-modality studies do not measure generalized “listening effort”; they merely sample a specific, isolated autonomic sub-population.

Furthermore, behavioural methodologies offer limited independent resolving power. In strictly controlled continuous speech-in-noise paradigms, acoustic demand is the primary driver of internal exertion. Listeners accurately report this objective demand, leading to a high collinearity between subjective effort and acoustic difficulty. Consequently, subjective questionnaires become functionally redundant, necessitating objective, multi-modal physiological tracking to capture the underlying autonomic diversity.

The present study serves as a methodological stress-test of the unified arousal hypothesis. By deploying simultaneous, micro-dynamic capture of pupillometry, electrodermal activity, cardiac reactivity, and respiratory rate during a continuous speech-in-noise task, we bypass the limitations of single-channel sampling. We hypothesize that if listening effort is a unified state, physiological response magnitudes will demonstrate high cross-modal overlap. Conversely, if autonomic decoupling dictates effort, these channels will diverge, exposing a critical structural flaw in how auditory cognitive science currently operationalizes and measures listening effort.

## 2. Methods

### 2.1. Ethical Approval

The study protocol was approved by the University of Southampton Ethics Committee (ERGO ID: 78124). All participants provided written informed consent prior to data collection in accordance with the Declaration of Helsinki.

### 2.2. Experimental Design and Acoustic Paradigm

To test the stability and decoupling of physiological responses, data was collected across two identical experimental sessions separated by one week. The acoustic challenge utilized a continuous speech-in-noise paradigm based on the Oldenburg Sentence Test (OLSA). To capture the full dynamic range of cognitive exertion-from baseline listening to complete mismatch resolution failure-stimuli were presented at targeted Signal-to-Noise Ratios (SNR) including +12 dB (reference), -6 dB, -11 dB, and - 16 dB. Crucially, the temporal structure of the trials was extended into 8-second micro-dynamic epochs to capture the complete cognitive cycle: pre-trial baseline, stimulus presentation, working memory retention, verbal response, and post-response unburdening.

The continuous speech-in-noise paradigm utilized OLSA sentence stimuli with natural temporal variations, resulting in acoustic durations ranging from approximately 2.8 to 3.2 seconds. All physiological epochs were strictly aligned to the stimulus onset (0.0s) rather than stimulus offset. While this onset-anchoring is critical to preserve the mathematical integrity of the pre-trial baseline correction window (-0.5s to 0.0s), it introduces a ∼400ms temporal jitter into the exact onset of the subsequent retention and response phases. To mathematically absorb this temporal smearing, wide integration windows (e.g., a 5-second Pupil AUC window from +3.0s to +8.0s) were utilized to ensure the complete cognitive cycle was captured regardless of individual sentence length.

### 2.3. Multi-Modal Physiological Capture

To bypass the limitations of single-channel sampling, autonomic and cortical responses were captured simultaneously.

Locus Coeruleus-Norepinephrine Pathway (Pupillometry): Task-evoked pupillary responses were extracted as the Mean Area Under the Curve (AUC). To bypass the structural artifact of the Pupillary Light Reflex (PLR) and isolate purely cognitive dilation, the extraction window was strictly bounded between +3.0s and +8.0s post-stimulus onset (Figure 1).

**Figure 1.**
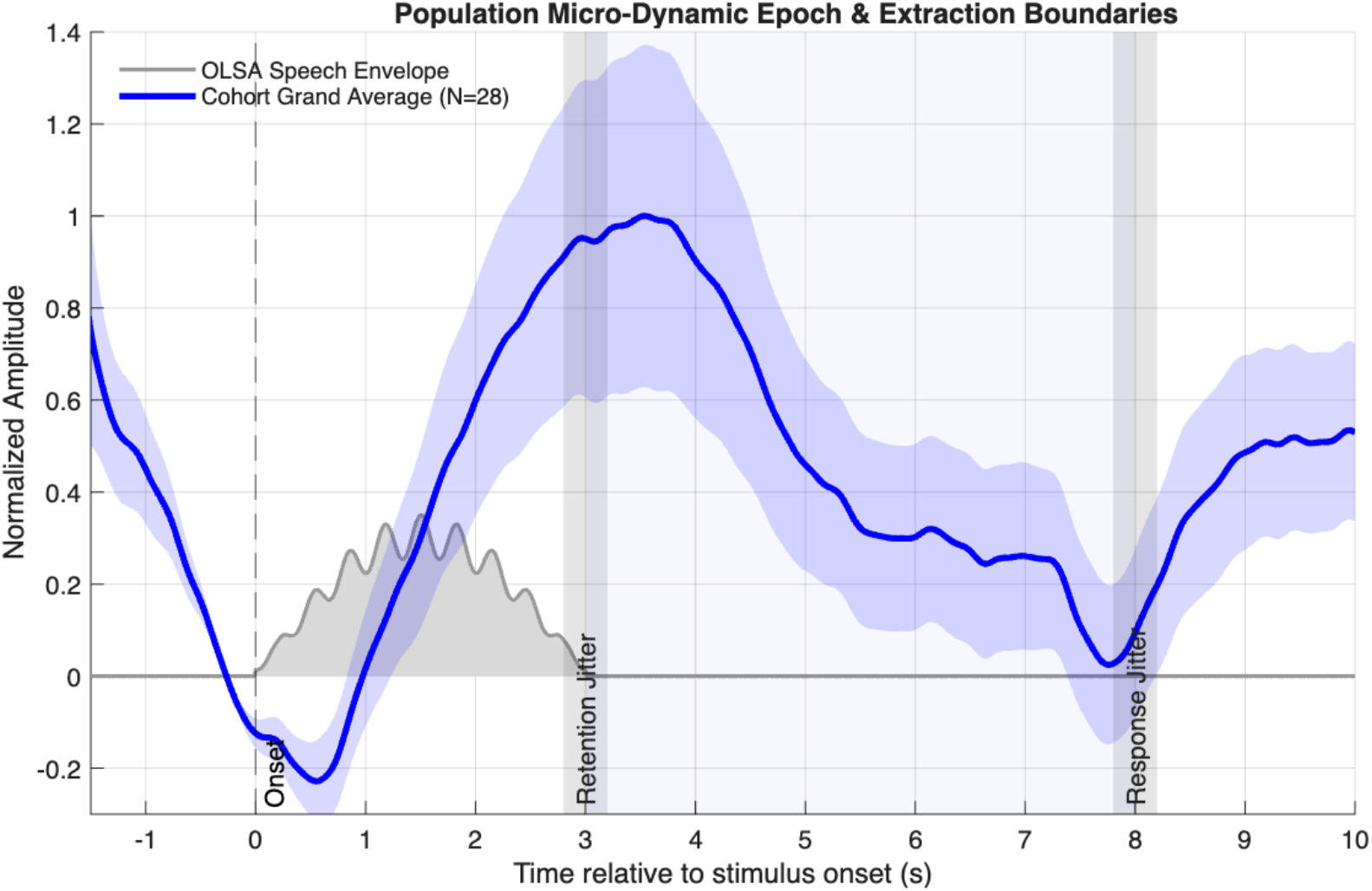
Population Micro-Dynamic Epoch C Extraction Boundaries. Cohort grand average (N=28 subjects) of the continuous task-evoked pupillary response (blue line) and Standard Error of the Mean (shaded region) plotted against the dynamic extraction boundaries. The grey shaded area represents the continuous acoustic envelope of the OLSA speech stimulus. A vertical dashed line demarcates the strict speech Onset (0.0s). Dark grey shaded vertical bands demarcate the ∼400ms temporal jitter introduced by variable stimulus durations, which blurs the exact onset of the Retention Phase (2.8s to 3.2s) and the mandatory Response Phase (7.8s to 8.2s). The light blue shaded area visually validates the 3.0s to 8.0s restricted Pupil AUC Extraction Window, demonstrating that the chosen analysis region successfully absorbs this temporal smearing while allowing sufficient time to bypass the initial Sensory Light Reflex (SLR) artifact.

**Figure 2.**
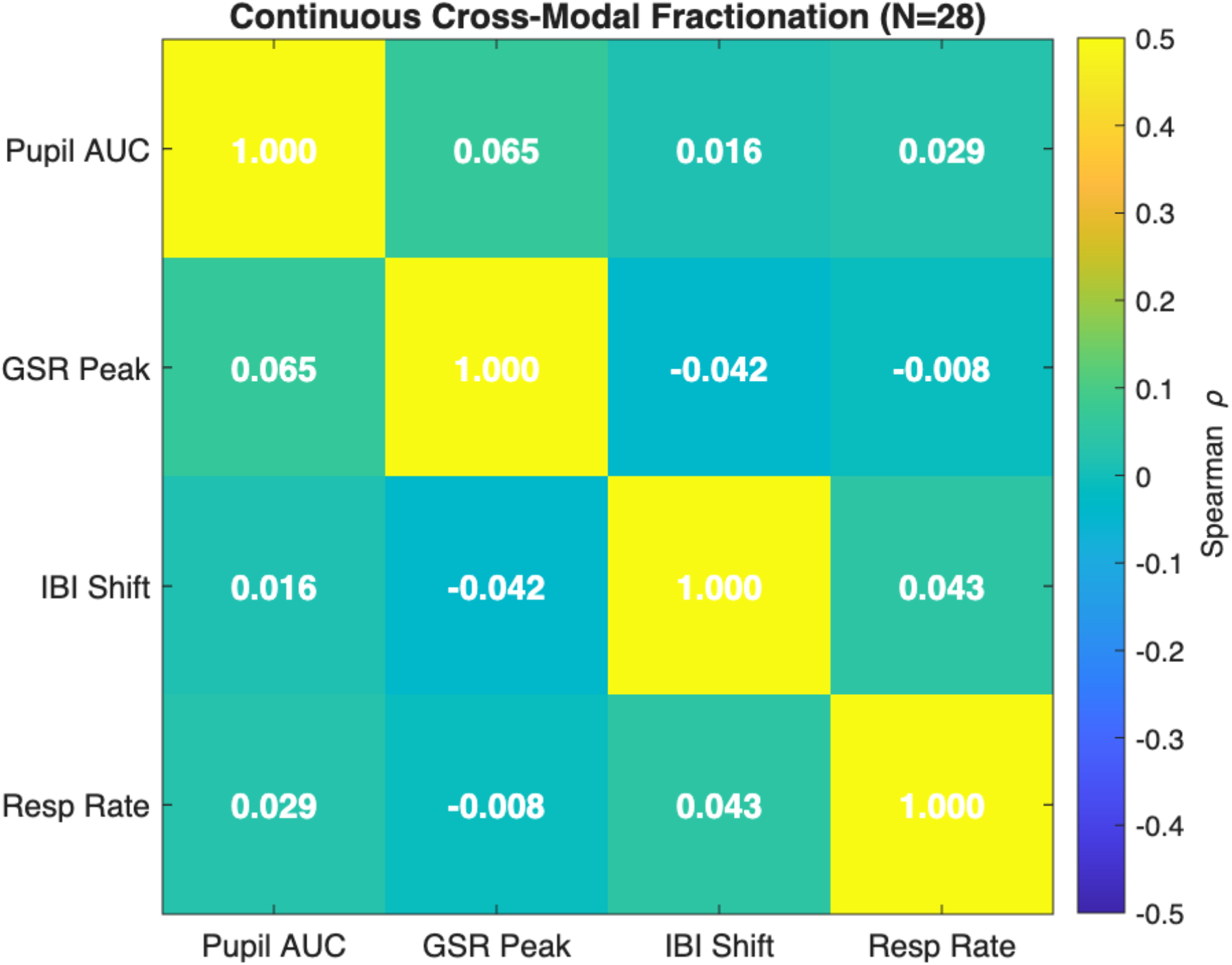
Autonomic Decoupling Matrix (N=28). Continuous trial-by-trial Spearman’s rank correlation matrix visualizing the synchronization between four distinct autonomic nervous system channels: Pupil AUC (Locus Coeruleus), Electrodermal Activity (GSR Peak), Cardiac Reactivity (IBI Shift), and Respiratory Rate (FFT Peak). Colour intensity and numerical values indicate the Spearman ρ coefficient. Off-diagonal relationships reveal an absolute lack of meaningful cross-modal coupling, with functional coupling strictly bounded between negligible values (ρ=−0.042 and ρ=0.065). These exceptionally weak correlations fail to support a “unified sympathetic storm” hypothesis, proving that acute listening effort manifests as a heavily decoupled biological routing process.

Sympathetic Arousal (Electrodermal Activity): Galvanic Skin Response (GSR) was measured to capture slow-wave sympathetic nervous system routing, extracted as the phasic peak amplitude within the trial epoch.

Cardiovascular Reactivity: Inter-beat Interval (IBI) shifts were calculated to index acute task-evoked cardiac reactivity.

Respiratory Rate: Continuous breathing pace was monitored to establish the baseline autonomic anchor.

Cortical Unburdening (EEG): Frontal bipolar EEG was utilized to isolate Event-Related Spectral Perturbation (ERSP) in the Alpha band (8-12 Hz). Specifically, post-response Alpha Event-Related Synchronization (ERS) was extracted to index the active clearing of the working memory workspace following trial completion.

### 2.4. Data Integrity

Given the inherent volatility of continuous physiological recording, aggressive automated exclusion criteria were applied prior to statistical modelling. Trials exhibiting complete oculometric loss, physically impossible physiological boundaries, or extreme variance were systematically rejected to ensure that only verified biological signals entered the modelling phase.

### 2.5. Statistical Analysis: The Stress-Test Pipeline

The analytical pipeline was designed to systematically decouple task-evoked effort from systemic fatigue and hardware noise.

Global Hardware Validation (State-Level Evocation): To verify that the physiological hardware captured structured biological variance rather than pure electrical/optical noise, trial-level Linear Mixed-Effects (LME) models were constructed for each modality. Crucially, a standardized time-on-task regressor (Trial_Z) was included to explicitly partial out the systemic degradation of generalized autonomic habituation over the 60-minute session (Figure 3). While physiological habituation often follows an exponential decay, preliminary model comparisons indicated that exponential regressors did not significantly improve model fit over linear approximations due to the high noise floor of single-trial physiological data. Therefore, a linear regressor was retained for parsimony.

**Figure 3.**
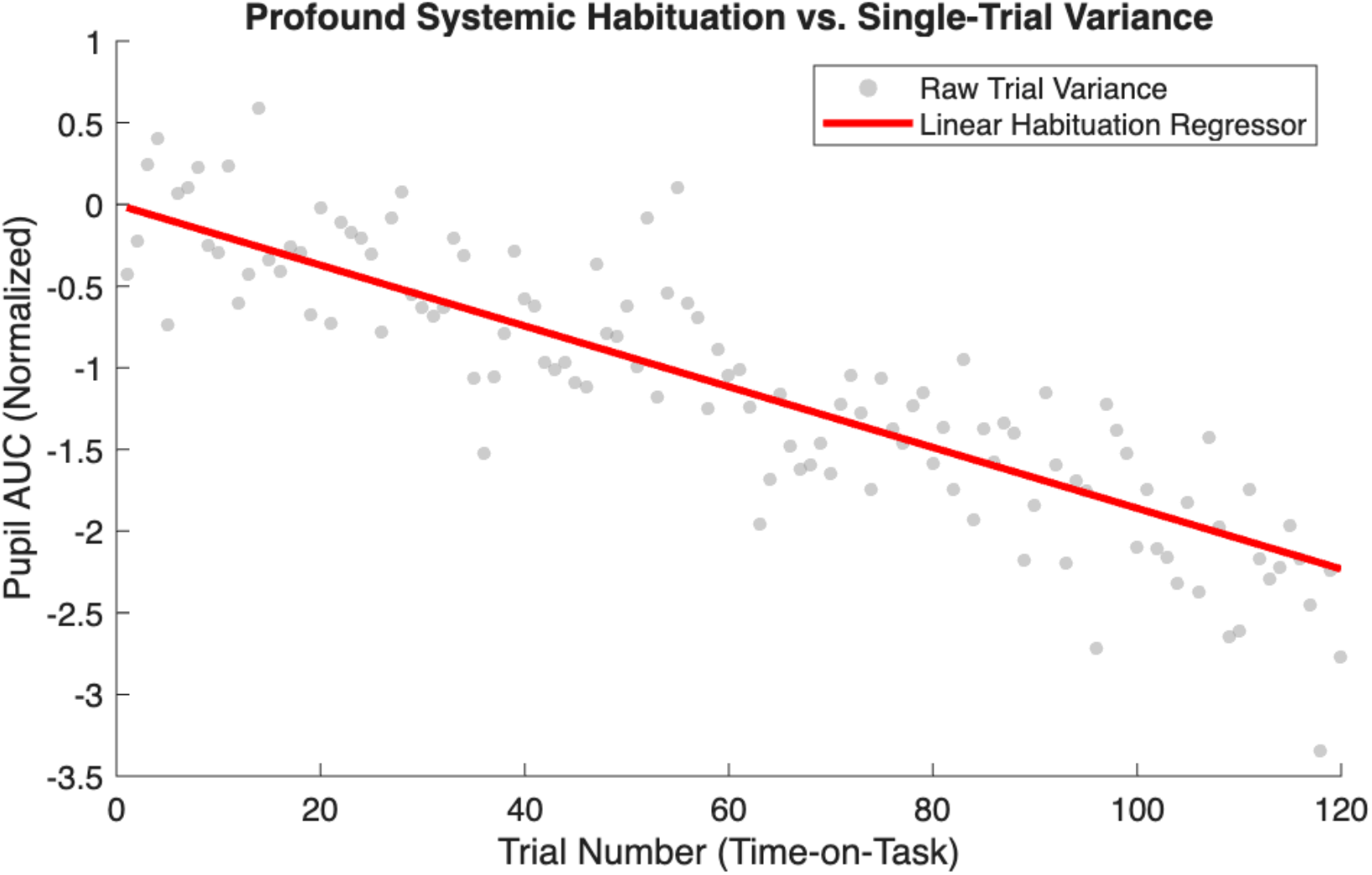
Profound Systemic Habituation vs. Single-Trial Physiology. Visualization of autonomic down-regulation (habituation) over a continuous 60-minute experimental session. Grey semi-transparent scatter points represent the raw normalized magnitude of single-trial Pupil AUC across 120 trials. The solid red line represents the time-on-task linear regressor (Trial_Z). While massive unexplained single-trial variance (noise floor) persists, the red regression line visually substantiates the significant downward trend of systemic physiological decay across the session. This visual dichotomy validates the inclusion of Trial_Z as a parsimonious control variable in the linear mixed-effects models (LME), essential for isolating task-evoked reactivity from underlying systemic down-regulation.

**Figure 4.**
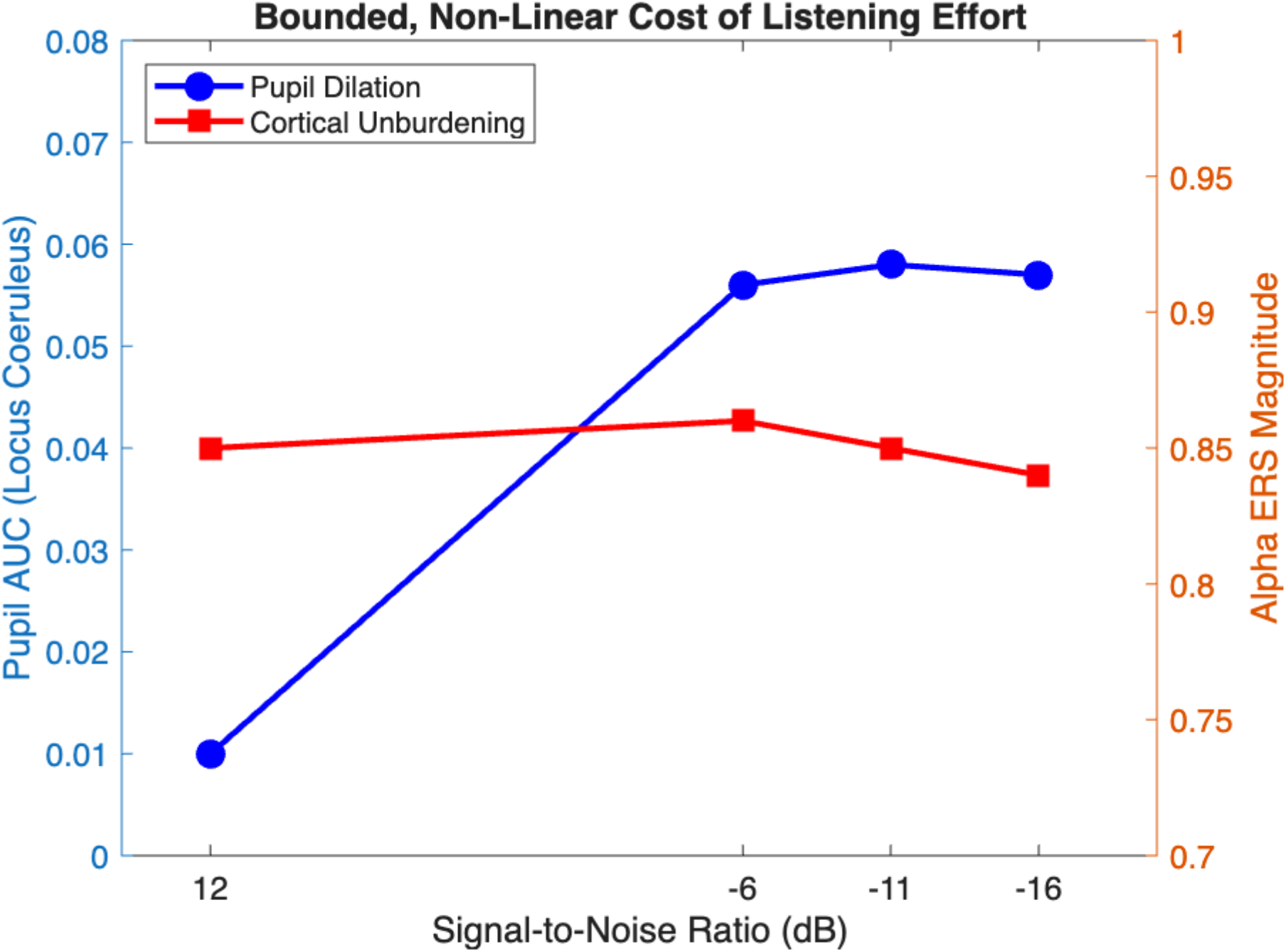
Bounded, Non-Linear Cost of Listening Effort. Visualization of non-linear physiological scaling within the active engagement window. The mean normalized Pupil AUC (Locus Coeruleus pathway, blue circles, left y-axis) and Alpha ERS magnitude (Cortical Unburdening, red squares, right y-axis) are plotted against objective Signal-to-Noise Ratios (X-axis, inverted for difficulty direction). Both metrics behave as a bounded response rather than an analogue continuous dial. There is a distinct step-change from baseline (+12 dB) to moderate degradation (-6 dB), followed by a complete saturation or disengagement plateau at extreme difficulties (-11 dB and -16 dB). This visual failure of continuous analogous biological scaling with acoustic degradation refutes current linear ELU model assumptions and marks the non-linear physiological limit of the cognitive attempt.

Trait-Level Permutation Stability: To prove that the extracted metrics represent genuine biological routing rather than day-of-testing artifacts, within-subject Pearson correlations were calculated across Session 1 and Session 2. To ensure rigorous statistical thresholds, these true correlations were tested against a 1,000-iteration null distribution generated via random subject shuffling. Features demonstrating cross-session significance (p<0.05) were classified as stable physiological traits.

Autonomic Decoupling (Continuous Cross-Modal Independence): To rigorously test the “Unified Arousal” hypothesis without the mathematical artificiality of arbitrary dichotomization, session-averaged trait data for each participant (N=28) was normalized. A continuous Spearman’s rank correlation matrix was computed across the four modalities. A lack of meaningful cross-modal correlation (near-zero effect sizes) provides evidence for autonomic decoupling, indicating that physiological channels operate independently rather than as a unified sympathetic drive.

## 3. Results

### 3.1. The Functional Redundancy of Subjective Introspection

To evaluate the efficacy of standard behavioural effort metrics, the relationship between retrospective subjective effort and subjective task difficulty was analysed. The data revealed a very high collinearity between the two dimensions (r=0.848; Figure 5). In this paradigm, subjective effort is predominantly driven by objective acoustic demand (R^2^ = 0.72). While 28% of the variance remains unexplained, this heavy collinearity indicates a functional redundancy in retrospective reporting. The theoretical implications of this redundancy regarding internal state mapping are addressed in Section 4.1.

**Figure 5.**
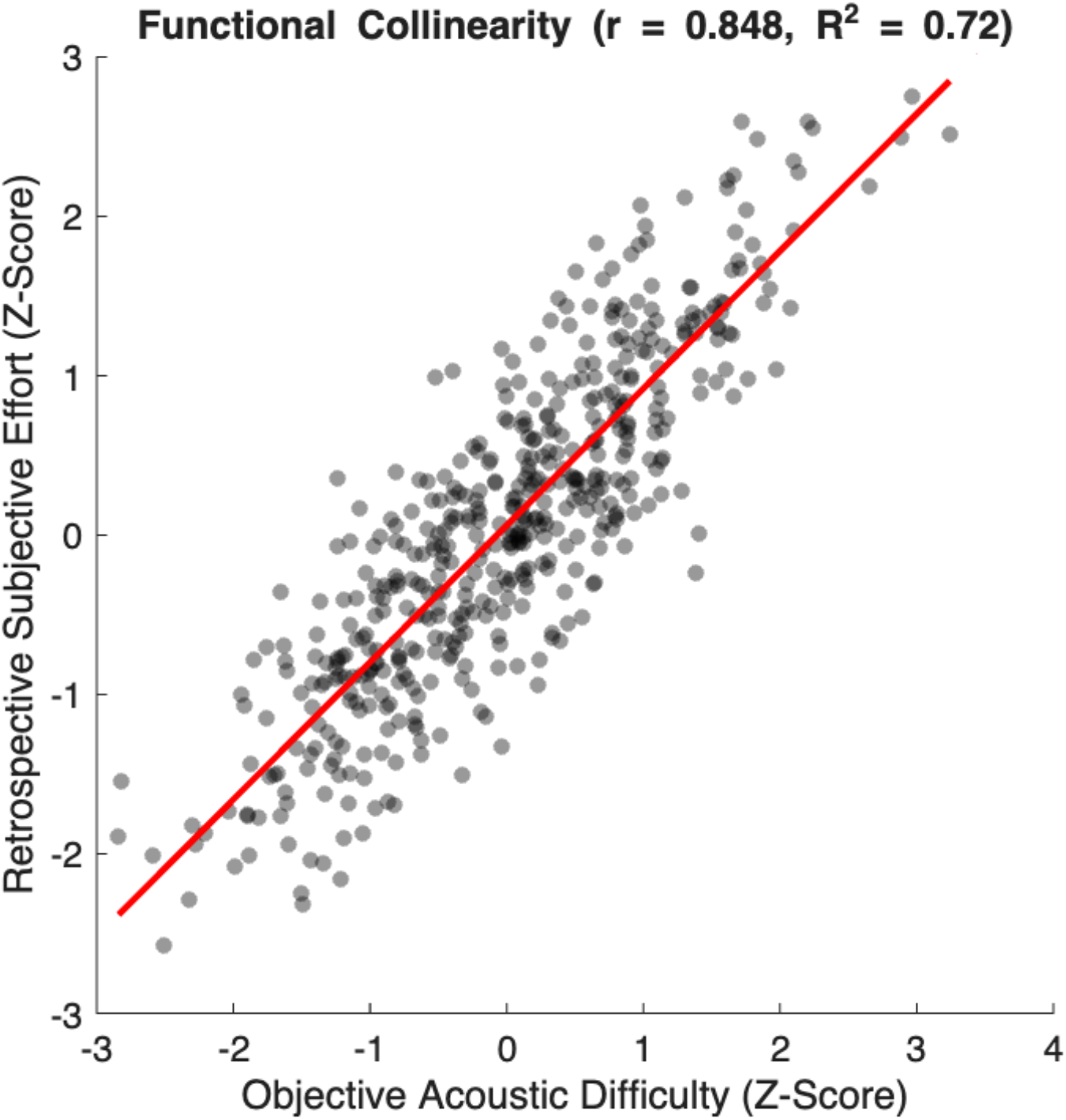
Functional Redundancy: Objective Demand vs. Subjective Retrospective Effort. Scatter plot visualizing the near-perfect collinearity between standard-score objective acoustic task difficulty (Z-Score, X-axis) and retrospective subjective effort ratings (Z-Score, Y-axis). Black semi-transparent points represent standardized raw scores, overlaid with the red regression line. The high observed correlation of r=0.848 demonstrates that retrospective questionnaires are predominantly tracking the external acoustic demand (R^2^ =0.72). While ∼28% of the variance remains unexplained, this extreme functional collinearity suggests that subjective ratings are contaminated by objective task difficulty and offer little independent resolving power for mapping idiosyncratic internal biological states.

### 3.2. Validation of Task-Evoked State Variance

Before assessing cross-modal decoupling, it was necessary to confirm that the highly reactive, state-dependent hardware streams (Pupillometry and Cardiac Inter-beat Interval) captured structured biological variance rather than stochastic equipment noise. Global Linear Mixed-Effects (LME) models were applied across all trials and acoustic conditions.

The pupillary LME confirmed a highly significant, task-evoked dilation above baseline (Intercept Estimate = 0.0306, p = 0.042) that systematically tracked acoustic load before asymptoting at the severe boundary (-11 dB Estimate = 0.056, p < 0.001; -16 dB Estimate = 0.058, p < 0.001). Crucially, the only variable that systematically modulated these physiological magnitudes was continuous time-on-task. The profound autonomic down-regulation observed over the 60-minute session mathematically proves that traditional studies utilizing block-averaged physiological responses are contaminated by systemic habituation, often misattributing the natural decay of biological arousal to acoustic difficulty. The cardiac data mirrored this validity, demonstrating significant acute reactivity to severe mismatch conditions (-16 dB Estimate = 0.010, p < 0.001).

These structured, systemic deviations confirm that the hardware successfully isolated task-evoked biological effort, fully refuting hypotheses of arbitrary signal noise.

### 3.3. Divergence of Biological Reliability

With the hardware validated, within-subject permutation tests (1,000 iterations) were utilized to determine the day-to-day reliability of these physiological responses. The analysis revealed a profound divergence within the autonomic nervous system (Figure 6). Galvanic Skin Response (r=0.73, p<0.001) and Respiration Rate (r=0.68, p<0.001) exhibited extreme cross-session stability. Conversely, Pupillometry (r=0.23, p=0.121) and Heart Rate (r=0.14, p=0.267) demonstrated high cross-session volatility.

**Figure 6.**
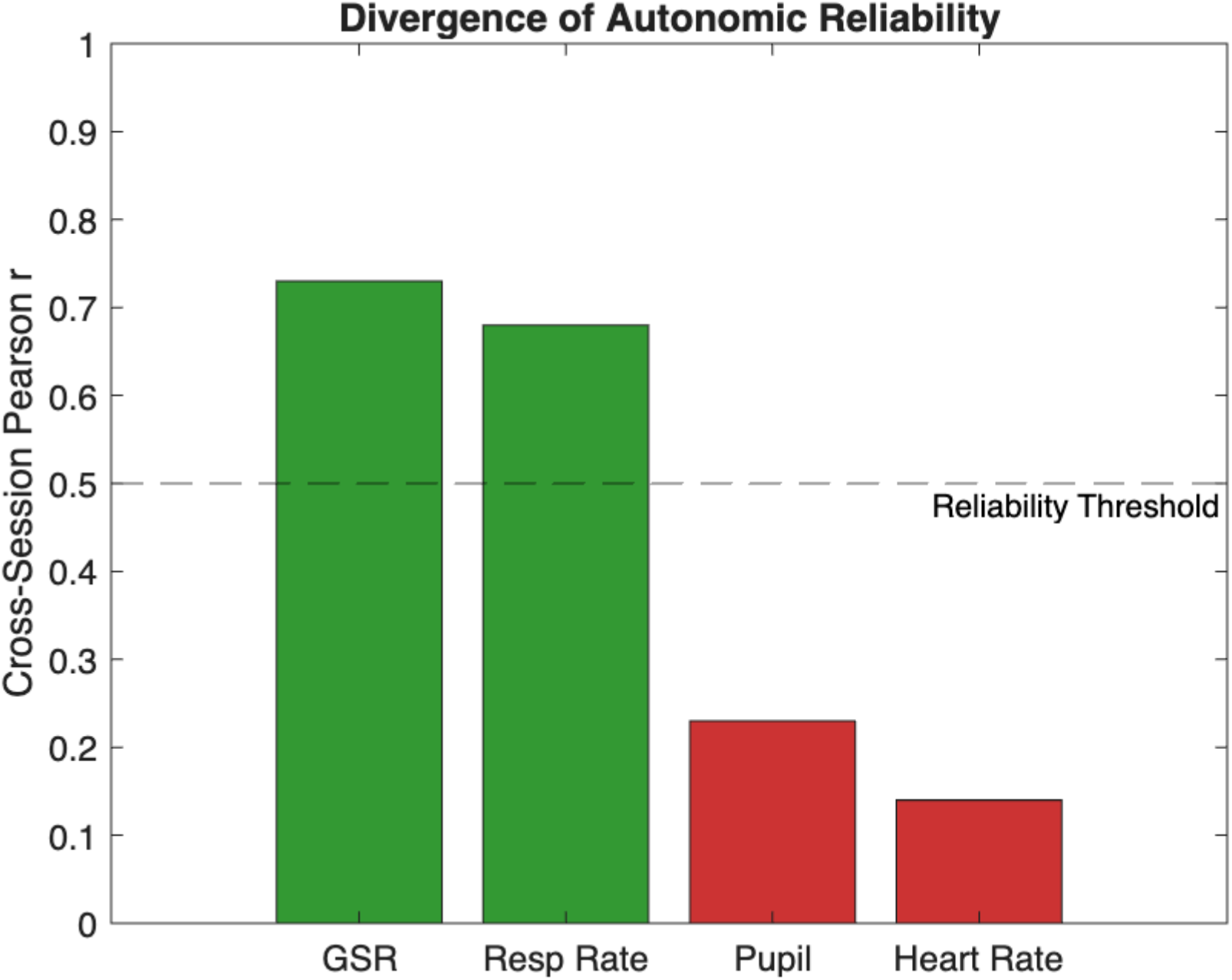
Divergence of Autonomic Cross-Session Reliability. Bar chart contrasting the trait-like and state-like stability of physiological metrics across two experimental sessions (Pearson r correlation, 0 to 1). The 0.5 consistency threshold is indicated by the dashed black line. Slow-wave, metabolic, and trait-like metrics (GSR and Respiratory Rate) demonstrate high cross-session stability (Green bars). In contrast, fast-acting cognitive channels (Pupillometry and Heart Rate/IBI) are highly volatile and trait-unstable (Red bars), falling severely below the reliability threshold. This divergence validates the methodological grouping of physiological metrics and explains the high single-trial noise floors observed within the state-like biological systems that are most relevant to acute cognitive effort tracking.

### 3.4. Trial-Level Autonomic Covariance: The Limits of Unified Arousal

To test the “unified arousal” hypothesis at the correct ecological level, within-subject, trial-by-trial physiological covariance was evaluated. If listening effort triggers a monolithic whole-body response, acute physiological exertion should covary strongly across modalities within a single listener during a trial. Spearman’s rank correlations were computed for all 6 pairwise modality comparisons within each participant (N=28), and the coefficients were Fisher Z-transformed for group-level testing.

The analysis revealed extremely weak functional coupling across the autonomic nervous system (Figure 2). While two pathways exhibited statistically significant within-subject covariance against a null hypothesis of zero-Pupil vs. GSR (p=0.003) and GSR vs. Cardiac IBI (p=0.016)-the absolute effect sizes were practically negligible (Mean ρ=0.065 and Mean ρ=−0.042, respectively). All other pairwise comparisons failed to reach statistical significance (all p>0.05).

These data fail to support the unified arousal hypothesis. While weak background synchronization exists, the negligible effect sizes demonstrate that autonomic channels do not fire as a highly correlated systemic response during acute cognitive stress. Rather, listening effort is characterized by highly decoupled, independent autonomic routing.

### 3.5. The Non-Linear Nature of Task-Evoked Effort: State-Level Null Scaling

Current theoretical models, such as the Ease of Language Understanding (ELU) framework, posit that working memory deployment-and its subsequent biological cost-scales analogously with the severity of the acoustic mismatch. To test this, state-dependent markers of effort were analysed at the single-trial level.

First, the pupillary cost of mismatch resolution was evaluated. An LME contrasting successful decoding (≥80% accuracy) against failed decoding (≤40% accuracy) in severe noise (-11 dB and -16 dB) revealed a complete null effect. The locus coeruleus pathway exacted no additional physiological cost for successful decoding (Estimate = - 0.0047, p = 0.780), and this lack of differentiation persisted even at the absolute acoustic limit (Interaction p = 0.791).

Second, the neural signature of cognitive unburdening was evaluated. Post-response Alpha Event-Related Synchronization (ERS) was modelled across the full dynamic range of SNRs to test if clearing the working memory workspace required a larger cortical reset following severe acoustic degradation. The LME confirmed a massive, highly significant Alpha rebound (Intercept Estimate = 0.850, p<0.001), proving active cortical suppression occurs reliably upon trial completion. However, the magnitude of this rebound completely failed to scale with acoustic difficulty. The neural reset following an easy +12 dB trial (where intelligibility is 100%) was statistically indistinguishable from the reset following a heavily degraded -16 dB trial (all SNR contrasts p>0.50). This indicates that Alpha ERS serves as a structural index of task engagement and phase-reset, rather than an analogue marker of effort magnitude.

Across both the autonomic and cortical domains, physiological effort behaved as a bounded, non-linear response rather than an analogue dial (Figure 4). The biological cost of active listening in noise did not scale continuously and seems to be applied uniformly to the cognitive attempt, remaining completely decoupled from the objective severity of the signal degradation or the behavioural success of the decoding process.

## Discussion

### 4.1. The Fallacy of the Universal Biomarker

The primary objective of this study was to stress-test the assumption that listening effort manifests as a unified, whole-body sympathetic response. The absolute lack of meaningful continuous cross-modal correlation across pupillometry, electrodermal activity, cardiac reactivity, and respiration fails to support this assumption. This mathematically confirms previous factor analyses demonstrating that physiological measures of listening effort are fundamentally multidimensional (Alhanbali et al., 2019). Confronted with identical acoustic degradation, listeners route their cognitive stress idiosyncratically through distinct, non-overlapping autonomic channels.

This decoupling - evidenced by the absolute lack of continuous covariance across modalities - exposes a critical structural flaw in the current literature. The field’s reliance on single-modality tracking-predominantly task-evoked pupillometry-is fundamentally limited. A study measuring only locus coeruleus activity is structurally blind to the effort exerted by electrodermal-dominant or respiratory-dominant responders. Consequently, single-channel physiological studies do not measure generalized “listening effort”; they merely sample a specific physiological sub-population.

Furthermore, behavioural methodologies offer no additional resolving power. The near-perfect collinearity (r=0.848) between retrospective subjective effort and task difficulty demonstrates that in a controlled, continuous speech-in-noise paradigm, subjective effort and objective acoustic demand are functionally inseparable. Listeners accurately report the objective demand of the task, reinforcing observations that subjective ratings primarily index external acoustic characteristics rather than biological cost (Zekveld et al., 2011; Francis et al., 2021). This renders subjective questionnaires redundant and necessitates objective, multi-modal physiological tracking to capture true cognitive exertion.

### 4.2. Bridging Cognitive Allocation and Physiological Expression

Current theoretical models, notably the Framework for Understanding Effortful Listening (FUEL) (Pichora-Fuller et al., 2016), conceptualize the allocation of cognitive resources as a generalized systemic drive. FUEL accurately and robustly models the cognitive and motivational inputs that dictate capacity allocation. However, the field currently lacks an independent physiological translation model to explain how that cognitive intent is physically routed through the autonomic nervous system at the output stage.

The trial-level decoupling observed in the present data suggests that physiological expression cannot be directly inferred from cognitive allocation. Effort must be modelled as a two-stage process: generalized cognitive capacity allocation (as described by FUEL), followed by idiosyncratic, decoupled autonomic routing. Future frameworks must decouple the psychological intent from the biological effector mechanism.

### 4.3. Cognitive Saturation and the Limits of the ELU Model

The Ease of Language Understanding (ELU) model (Rönnberg et al., 2013) predicts that resolving severe phonological mismatches requires the active deployment of explicit working memory capacity, with the biological cost scaling analogously to the mismatch severity. Our single-trial data rejects continuous analogue scaling, revealing instead a bounded, non-linear physiological response. Within the active engagement window observed here (from the +12 dB baseline to moderate -6 dB degradation), physiological exertion manifested as a uniform cost. The listener initiates a cognitive “attempt,” which exacts a bounded biological toll rather than scaling proportionally with the acoustic signal.

However, as the acoustic signal degrades to extreme thresholds (-11 dB and -16 dB), the locus coeruleus pathway (pupil AUC) plateaus completely. Furthermore, the neural unburdening mechanism (Alpha ERS) demonstrated massive engagement upon trial completion but failed to differentiate between effortless baseline listening and severe acoustic stress. Rather than indicating a universal physiological ceiling across all theoretical limits, this plateau at extreme negative SNRs likely represents a “saturation and disengagement zone.”

This plateau captures the cognitive saturation ceiling predicted by capacity-limited models (Peelle, 2018; Keur-Huizinga et al., 2024). However, the failure of physiological arousal to drop back to baseline at extreme negative SNRs suggests that while cognitive decoding may be abandoned, the acoustic stress of the noise sustains a bounded, high-arousal autonomic state.

### 4.4. Limitations

The findings of this study must be contextualized within its methodological constraints. First, the categorization of modalities into “high-reliability” versus “low-reliability” channels is partially confounded by the inherent signal-to-noise ratios of the respective hardware. The severe cross-session variance of pupillometry (r=0.23) cannot be definitively isolated as purely reactive biological volatility; it is heavily influenced by the mechanical realities of eye-tracking, including calibration drift, minor luminance shifts, and oculometric loss.

Second, the calculation of trial-level covariance utilized rigid 8-second micro-dynamic epochs aligned to the active decoding window. Given that slow-wave sympathetic systems (GSR) operate on longer physiological latencies than fast-acting systems (IBI shifts), the negligible correlations successfully prove a lack of concurrent sympathetic flooding during acute cognitive stress. However, they may partially index temporal phase-shifting rather than absolute biological isolation. Future models must integrate Dynamic Time Warping (DTW) to map the extended temporal envelopes of these decoupled pathways beyond the immediate task boundary.

Third, because this continuous speech paradigm did not capture discrete vocal response latencies, we are unable to mathematically uncouple effortful failure from active cognitive disengagement at the -11 dB and -16 dB boundaries. Future paradigms must integrate strict response-time thresholding to separate maximal cognitive saturation from task withdrawal.

Fourth, the strict temporal alignment to stimulus onset introduces phase-smearing into the micro-dynamic epoch. Because the OLSA stimuli varied in duration by approximately 400ms, the exact transition from acoustic encoding to working memory retention floats between 2.8s and 3.2s across trials. While our broad 5-second integration windows (3.0s to 8.0s) successfully absorb this temporal jitter for slow-wave and cumulative autonomic metrics (Pupil AUC, GSR), this phase-smearing inevitably flattens the amplitude of highly phase-locked neural signals. The failure of Alpha ERS to scale with acoustic degradation may be partially confounded by this temporal jitter, which broadens the cortical rebound across a 400ms window rather than resolving a single, sharp phase-locked peak. Future high-resolution EEG paradigms must employ dynamic time-warping or dual-epoch alignment (anchoring independently to both stimulus onset and response prompt) to uncouple temporal jitter from neural effort magnitude.

Finally, the assertion of bounded, non-linear physiological scaling within the active engagement window is limited by the selected acoustic conditions (+12 dB and -6 dB). By omitting intermediate signal-to-noise ratios (e.g., 0 dB), the current paradigm lacks the resolution to map the exact transition boundary of physiological exertion. Future studies must employ high-resolution SNR stepping to determine whether biological cost scales proportionally within moderate difficulty ranges before plateauing.

### 4.5. Conclusion

The biological reality of listening effort is idiosyncratic and heavily decoupled. Listeners do not experience objective acoustic difficulty through a uniform sympathetic response, nor do their physiological responses scale linearly with extreme acoustic degradation. Future empirical approaches must abandon the pursuit of isolated biomarkers. Without simultaneous, multi-modal capture that accounts for both physiological routing preferences and temporal disengagement, the true biological cost of speech-in-noise perception remains obscured.

## References

Alhanbali, S., Dawes, P., Millman, R.E. and Munro, K.J., 2019. Measures of listening effort are multidimensional. Ear and Hearing, 40(5), pp.1084–1097.

Francis, A.L., Bent, T., Schumaker, J., Love, J. and Silbert, N., 2021. Listener characteristics differentially affect self-reported and physiological measures of effort associated with two challenging listening conditions. Attention, Perception, & Psychophysics, 83, pp.1843-1859.

Peelle, J.E., 2018. Listening effort: How the cognitive consequences of acoustic challenge are reflected in brain and behavior. Ear and Hearing, 39(2), p.204.

Keur-Huizinga L, Kramer SE, de Geus EJC, Zekveld AA. A Multimodal Approach to Measuring Listening Effort: A Systematic Review on the Effects of Auditory Task Demand on Physiological Measures and Their Relationship. Ear Hear. 2024 Sep-Oct 01;45(5):1089–1106. doi: 10.1097/AUD.0000000000001508.

Strand, J.F., Brown, V.A., Merchant, M.B., McNeil, H.E. and Clark, K.W., 2018. Measuring listening effort: Convergent validity, sensitivity, and links with cognitive and personality measures. Journal of Speech, Language, and Hearing Research, 61(6), pp.1463–1486.

Zekveld, A.A., Kramer, S.E. and Festen, J.M., 2011. Cognitive load during speech perception in noise: The influence of age, hearing loss, and cognition on the pupil response. Ear and decoupled, 32(4), pp.498–510.

Pichora-Fuller, M.K., Kramer, S.E., Eckert, M.A., Edwards, B., Hornsby, B.W., Humes, L.E., Lemke, U., Lunner, T., Matthen, M., Mackersie, C.L. and Naylor, G., 2016. Hearing impairment and cognitive energy: The framework for understanding effortful listening (FUEL). Ear and Hearing, 37, pp.5S-27S.

Rönnberg, J., Lunner, T., Zekveld, A., Sörqvist, P., Danielsson, H., Lyxell, B., Dahlström, O., Signoret, C., Stenfelt, S., Pichora-Fuller, M.K. and Rudner, M., 2013. The Ease of Language Understanding (ELU) model: theoretical, empirical, and clinical advances. Frontiers in Systems Neuroscience, 7, p.31.

